# The nanopore sequencing of a Chinese rhesus macaque revealed patterns of methylation, recombination, and selection for structural variations

**DOI:** 10.1101/2022.10.21.513306

**Authors:** Jianhai Chen, Jie Zhong, Xuefei He, Ivan Jakovlić, Yong Zhang, Hao Yang, Younan Chen, Guang Yang, Chuanzhu Fan, Bairong Shen

## Abstract

Rhesus macaques (Macaca mulatta) are the most extensively studied nonhuman primate species for human biomedical modeling. However, little is known about the biological pattern of genome-wide structural variations (SVs) and the evolutionary forces underlying SVs. Here, we conducted genomic sequencing and analyses based on Nanopore long reads and Illumina short reads technology. We called SVs between the two subspecies (China vs. India), using three methods of assembly-based and long-reads-based algorithms. Interestingly, we found significantly more SVs in X-chromosome than in autosomes, consistent with the expectation of the faster-X divergence at the subspecies level. With the fine-scale methylation frequencies and recombination rates, we found duplications with significantly lower methylation frequencies while higher recombination rates than other types of SVs, suggesting a higher level of transcriptional and evolutionary potential for duplications than for other SVs types. A genome-wide scan of selective sweep revealed that over 3% of SVs are under positive selection. Moreover, X chromosome showed significantly higher number of positively selected SVs than do autosomes, suggesting the “faster-X effect” of SVs. Our study revealed a different evolutionary importance for duplications compared with other SVs forms. We also revealed the “faster-X effect” of SVs, which could provide raw material upon which positive selection can further play.

## Introduction

Rhesus macaque (Macaca mulatta) has been extensively studied in biomedical field for human diseases [2]. This nonhuman primate model has greatly expanded and deepened our understanding on both infectious and genetic diseases, such as AIDS [3, 4], Ebola [5], SARS-CoV-2 [6, 7], autism [8-10]. Parkinson’s disease [11, 12], cataract [13, 14], cardiovascular diseases [15], infertility [16, 17], etc. This popularity is probably due to its biological, evolutionary, and ecological features, including adaptive flexibility, widespread distribution, population abundance, genetic closeness to human than other model species (i.e., mouse, fruit fly, and zebrafish), and highly diverse genomic variants [18, 19]. As one of the most evolutionarily and ecologically successful nonhuman primates [20, 21]. It has now occupied the widest natural habitats, extending from Pakistan and Afghanistan in the west across South Asia to Southeast Asia and the eastern coast of China [22]. Taxonomically, the best characterized rhesus macaques are from two well-differentiated subspecies: the Indian subspecies and the Chinese subspecies, which diverged at ∼162,000 years ago [23].

Genomic SNPs derived from the next-generation sequencing (NGS) data have greatly promoted our understanding on evolutionary history and biomedical relevance of the macaque species [24-26]. However, it is still lacking the SVs-oriented studied to demystify basic and evolutionary patterns of SVs. In this study, we are particularly interested in whether genomic SVs could be shaped by chromosomal distribution, methylation frequencies, recombination rates, and evolutionary forces. The distribution of SVs and the related evolutionary forces could provide a new empirical evidence for the “faster-X effect”, an evolutionary pattern caused by differences between autosomes and hemizygous sex chromosomes [27-30]. The relationship between SVs and DNA methylation is a still unclear, which could be addressed with the nanopore long-read sequencing due to its direct signals of epigenome-wide methylation [31, 32]. In addition, NGS data could be incorporated to infer the fine-scale recombination rates [33], which may address issues on the recombination bias of SVs. Thus, by combining both NGS data and nanopore long-read sequences, we could achieve a more balanced and nuanced perspective on the genomic features of SVs using rhesus macaque as a model.

As a prerequisite for in-depth genetic studies, the genome assemblies of rhesus macaques placed an indispensable role in decoding evolutionary and biomedical questions. Before the advent of long-read sequencing, the next-generation sequencing (NGS) data were utilized to construct the draft genome of the Indian rhesus macaque (rheMac8) [2]. A decade later, a long-read assembly for the Chinese rhesus macaque was established using the Pacific Biosciences’ (PacBio) single-molecule real-time (SMRT) and NGS data [34]. With the metric N50, the shortest sequence length needed to cover 50% of the genome [35], this reference genome (rheMacS) achieved an elevated level of continuity (8.19Mb of contig N50 and 13.64Mb of scaffold N50) than rheMac8 [34]. Three years later, also using the PacBio long-read and NGS data, the reference of the Indian rhesus macaque (Mmul_10) was improved significantly (∼120-fold) in sequence contiguity (46Mb of contig N50) relative to rheMac8 [19]. In recent years, another big step forward for *de novo* genome assembly comes from the development of the Nanopore long-read sequencing with its cost-effective advantage [36, 37], which improved the genome references for multiple species, such as humans [38-41], mammals [42-44], birds [45, 46], plants [47-49], fishes [50, 51], fruit flies [52], etc. However, there is still no nanopore derived reference assembly and related analyses (such as methylation, recombination rate, etc.) for rhesus macaques. Here, we initially provided a *de novo* assembly for the genome of a male Chinese rhesus macaque subspecies (CR2) with Nanopore long-read and Illumina short-read data. We achieved scaffold N50 of 149.74 Mb, slightly higher than previously reported genome of both India (mm10, 148 Mb) and Chinese rhesus macaque (rheMacS, 145 Mb). Using this assembly, nanopore long reads and NGS data, we identified SVs, estimated methylation frequencies, inferred recombination rates, and scanned selective sweep signals. We found an excess distribution of long SVs (1Kb) in X chromosome than in autosomes. We further revealed lower methylation frequencies but higher recombination rates for duplications than for other SVs (deletions, insertions, and inversions). We finally uncovered that some SVs are within the regions under selective sweep, suggesting the positive Darwinian selection would impact the evolutionary fate of these SVs at the population level. We also found that the X chromosome contributes more to the positively selected SVs-involved genes, suggesting the “faster-X effect” at subspecies level. Our analyses provide insights into the patterns of SVs and their diverse levels of methylation, recombination, and selective forces.

## Results

### The sequencing and *de novo* assembling

To determine the population identity of our focal sample of a Chinese rhesus macaque (CR2), we obtained 292,478,835,000 base pairs of Illumina short-read data (95.27x). Subsequently, we performed population genetic analyses based on totally 27 samples, including the newly sequenced and publicly available NGS data for Chinese and Indian rhesus macaques. We confirmed the subspecies ancestry of CR2 with the method the principal component analysis (PCA). We found that CR2 was genetically within the cluster of Chinese rhesus population instead of the Indian population (Figure 1A). This result supported CR2 as an acceptable representative of Chinese rhesus macaque for further sequencing and subsequent analyses.

**Figure 1.**
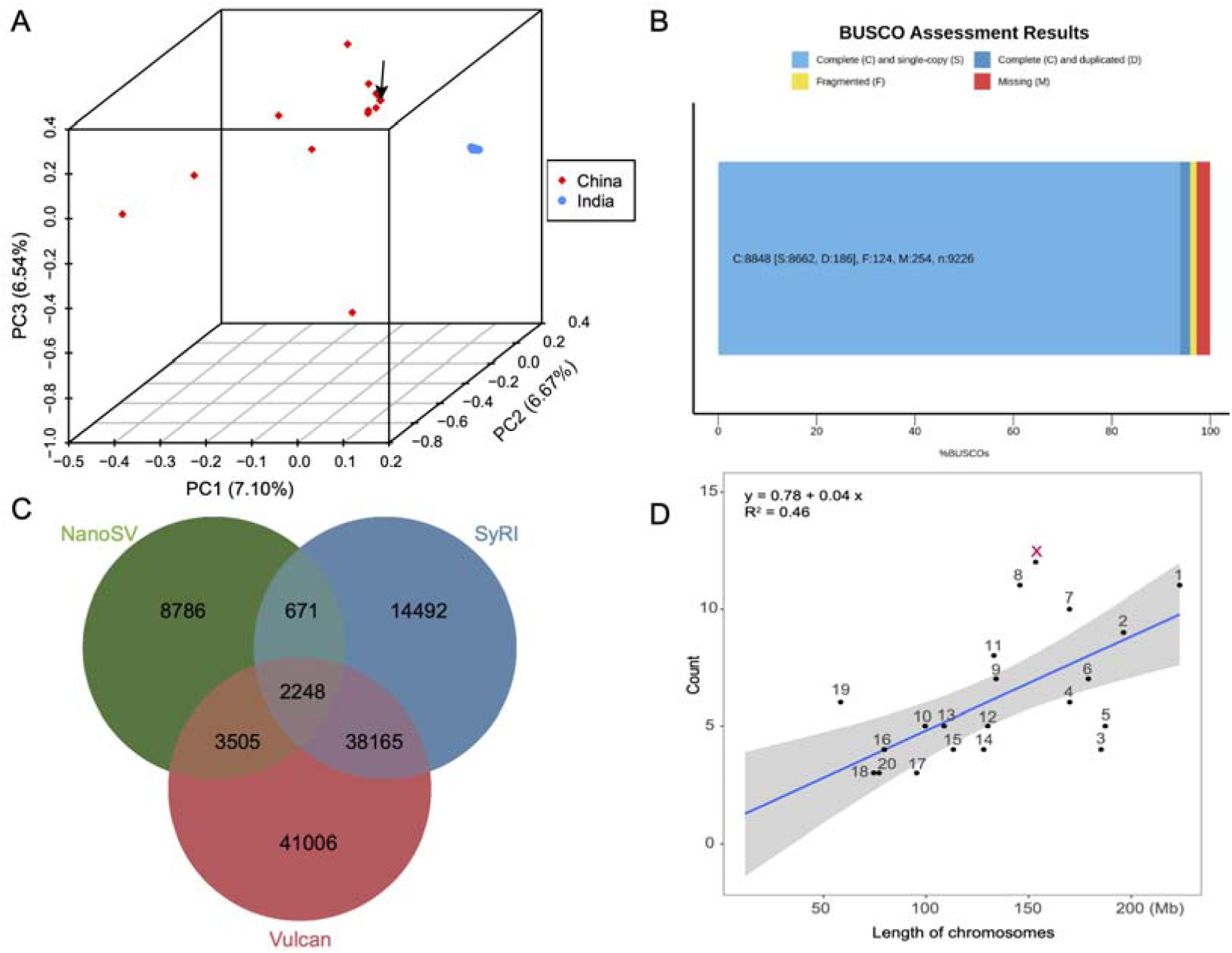
(A) The population identify for the newly sequenced sample based on PCA analysis. The arrow shows the sample sequenced in this study. (B) The completeness with Benchmarking Universal Single-Copy Orthologs (BUSCO) for the final reference genome. (C) The numbers of unique and sharing SVs for three methods of SVs identification (Vulcan, NanoSV, and SyRI). (D) The number of long SVs (1Kb) within chromosomes. The linear regression was based on the autosomes (“lm” function in R platform, p < 0.001).

For the purpose of SVs calling using the “assembly-based” method [53], we initially conducted *de novo* assembly of CR2 based on both NGS and long-read data. In total, we obtained 98,513,621,108 base pairs of Nanopore long-reads, with read N50 length 49.91 Kb and average read length 26 Kb from ONT-PromethION platform. Based on the estimation of GCE v1.0.2 with kmer 17 [54], the genome size of Chinese rhesus macaque is ∼□3.12□Gb. Thus, we achieved ∼32x genome coverage of long reads. A hybrid and cost-effective methodology was then performed using these long reads and ∼95x short reads. Previous reports have revealed that a hybrid assembly would need over 50x short-read coverage and over 30x long-read coverage [55, 56]. Thus, the data volume in this study is sufficient for *de novo* assembly. The major steps of genome assembling included reads cleaning (NECAT v0.0.1 and fastp), *de novo* assembling (NECAT v0.0.1 and NextDenovo v2.4.0), reference-guided chromosome assignment (RagTag v2.1), long and short reads correction (Pilon v1.24, Racon v1.4.3, Inspector v1.0.2, and Tigmint v1.2.6).

We found 14.87 times more contigs with NECAT than with NextDenovo (3376 vs. 227), probably because NECAT captured more smaller contigs than NextDenovo. In addition, the total length of contigs from NECAT is longer than that from NextDenovo (3.13Gb vs. 2.85Gb), suggesting a higher yield but more fragmented genome from NECAT than from NextDenovo. In addition, NextDenovo achieved a higher contig N50 than previously reported two references (65.99Mb vs. 46.61Mb and 8.19Mb). Thus, we jointly utilized long contigs from NextDenovo and shorter contigs from NECAT following the RagTag “scaffold” function and reduced the number of contigs (2182) and meanwhile maintained higher genome size (3.13 Gb).

Because of no available optical mapping or Hi-C data for our sample, we took a reference-guided method (rheMacS) to assign, order, and orient some contigs into the chromosomes [34]. We reduced the number of contigs into 1,479, lower than the previous two references (Table 1). After applying both long and short reads correction, we obtained a chromosome-level genome reference for CR2 (Table 1 and Supplementary figure 1). The full-length nanopore RNAseq data, which were sequenced based on pooled tissues of heart, liver, whole brain, intestine, testis, muscle, and pancreas, were used for gene annotation.

**Table 1.** The QUAST statistics for previously reported two references (rheMacS and Mmul_10, PacBio long reads) and the new reference (CR2, Nanopore long reads).

| Statistics | rheMacS | Mmul_10 | CR2 |
| --- | --- | --- | --- |
| Data | PacBio Sequel | RSII | Nanopore,<br>Illumina |
| Tools | FALCON | HGAP4_SMRT_Link | B* |
| # contigs (>1Kb) | 4,743 | 3,167 | 2,182 |
| # scaffolds (>1Kb) | 1,881 | 2,924 | 1,479 |
| Largest contig | 45,429,312 | 98,978,568 | 91,298,995 |
| Total length | 3,037,157,472 | 2,971,329,548 | 3,125,302,881 |
| GC (%) | 41.03 | 41.00 | 41.01 |
| Contig N50 | 8,187,455 | 46,608,966 | 65,986,596 |
| Scaffold N50 | 148,403,059 | 145,679,320 | 149,742,375 |
| # N's per 100 kbp | 179.86 | 1159.04 | 2.69 |

**Supplementary figure 1.**
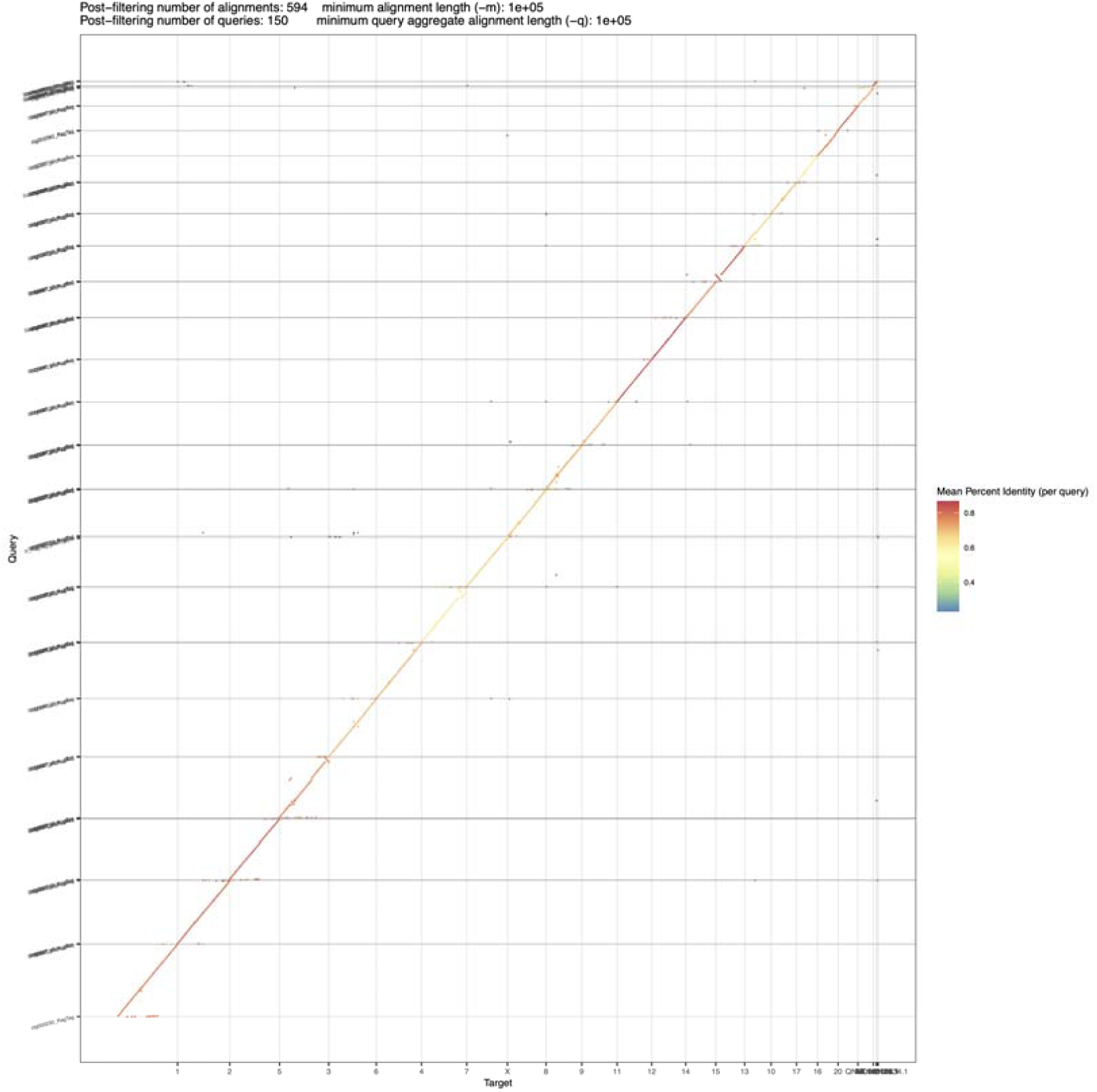
The whole genome alignment between CR2 and reference genome of an Indian rhesus macaque (mm10) using minimap2 (-xasm10). The dotplot visualization was based on a R package dotPlotly. The vertical axis shows the CR2 scaffolds while the horizontal axis exhibits the chromosomes of mm10.

Previously, two genomic references for the Chinese (rheMacS) and Indian rhesus macaque subspecies (Mmul_10 or mm10) were based on mainly the PacBio long reads [34]. In this study, the long reads of CR2 were obtained from Nanopore technology (R9.4), which offered an advantage of methylation information as well as the long reads. The comparison among summary statistics of three references revealed that CR2 has a slightly improved N50 values and lower counts for contigs and scaffolds, relative to both rheMacS and Mmul_10 (Table 1). The BUSCO evaluation [57], which is based on 9226 near-universal single-copy genes in mammalian species (odb10), revealed that the compete BUSCOs is 95.9% (Figure 1B), slightly higher than the previously reported percentage of a Chinese rhesus macaque (rheMacS) [34].

### Genes missing in the mm10 assembly were uncovered

Mouse, rat, and macaque are among the most widely used animal models for human medical issues. Despite closer evolutionary relationship between human and macaque, mouse and rat are oftentimes better choice for human transgenic studies due to not only the operation convenience of mouse and rat but also the missing of orthologous genes in current macaque genome. We tried to understand how many human orthologous genes are present in mouse but absent in macaque. Based on Ensembl annotation (v105) of comparative species, we retrieved “one to one “ orthologous genes between human, mouse, rat, and macaque and obtained human orthologous genes present in rodent species (mouse and rat) but absent in macaque. After mapping these human genes back to mm10 genome reference, we obtained 89 genes without reliable mapping coordinates of orthologous genes in macaque, using even the most relaxed threshold (Evalue 0.001 in TBLASTN). These results suggest that these genes may be either bona fide evolutionarily lost genes or hidden genes due to annotation error in genome reference of mm10.

To identify the potential hidden genes, we firstly produced a reference-guided assembly of transcripts based on Nanopore direct RNA sequencing data from a pool of seven tissues and then used these transcripts as the reference to retrieve the 89 potential hidden genes. Notably, 75.28% of these genes (67 out 89) were mapped to transcripts with reliable signals (E-value < 0.001) and 65 genes showed E-value less than 10^−10^. The coding sequences of these genes showed a slightly but not significantly higher median value of GC contents (51.93%) relative to genomic background level (51.45%). The “absence” of these “hidden genes” could be due to the previously neglected complex transcripts using only NGS data [58], suggesting the importance of long-read RNAseq data in decoding a more complete spectrum of transcriptome.

### Excess number of SVs was found in X chromosome

Genomic SVs between the Chinese and Indian subspecies was identified jointly with the read-based and assembly-based strategies [59] of three methods, NanoSV [60] and Vulcan [61], and SyRI [53]. The two methods based on long-read data could counteract the effects of assembly errors, thereby reducing the bias caused by only the assembly-based strategy. In addition, since the samples from the Chinese and Indian subspecies (CR2 vs. mm10) were compared, the SVs results could largely represent the subspecies-level divergence.

Specifically, Vulcan (41006) showed a more sensitive calling of SVs than both NanoSV (8786) and SyRI (14492). Interestingly, Vulcan and SyRI identified more SVs than Vulcan and NanoSV did, although Vulcan and NanoSV are reads-based methods while SyRI is an assembly-based method. Using a rigorous criterion (>80% of reciprocal overlapping region for each SV), 2248 congruent SVs with were found among all three methods (Figure 1C and Supplementary table 2). It is worth noting that some previous studies preferred to use the shared SVs obtained from different algorithms as the bona fide call-set of SVs [62]. For the specific purposes of this study, we used a reconciled method by treating the congruent SVs as the lower bound while the independent SVs from three methods as the upper bound of the bona fide call-set.

Based on the congruent 2248 SVs, we analyzed the chromosomal distribution preference of SVs. The percentage of X-chromosomal SVs is not the highest (4.54%) relative to some autosomes (7.87% for chromosome 1, 7.12% for chromosome 19, 5.83% for chromosome 11, etc.). However, after adjusting the effective population size ratio between X and autosomes (3/4) and considering the physical length of chromosomes, we found an excess of X-chromosome SVs relative to autosomal SVs (0.89/Mb vs. 0.80/Mb). When focusing on the long SVs over 1000bp, we found that X chromosome was the most extreme outlier of the linear pattern that shapes autosomes (Figure 1D, p < 0.001). This pattern indicates the “faster-X divergence” in principle [63], which is consistent with a previous observation in other mammalian species based on segmental duplication identified using only SyRI [27].

### The genome-wide methylation and structural variations (SVs)

The Nanopore sequencing can capture the genome-wide signals of methylation [64], which provides us a unique chance to understand the epigenetic pattern at a more diverse scale for genomic elements. The genome-wide methylation frequencies showed a similar distribution pattern across autosomes while a slightly flatter distribution on X and Y chromosome, suggesting different methylation patterns between homozygotes and hemizygotes (Supplementary figure 2 and Figure 2a). Median methylation values were found to be significantly different between autosomes (0.812), X (0.800), Y (0.778), and mitochondrial genome (0.032) (Figure 3a, Wilcoxon test, p < 0.001). The distribution shape of methylation frequencies in mitochondrial genome revealed a single small value peak (0.015), which is in sharply contrast to the peaks of larger values for nuclear chromosomes (>0.8, Figure 2B). Thus, our nanopore data supported the low methylation level in mitochondrial genome.

**Supplementary figure 2.**
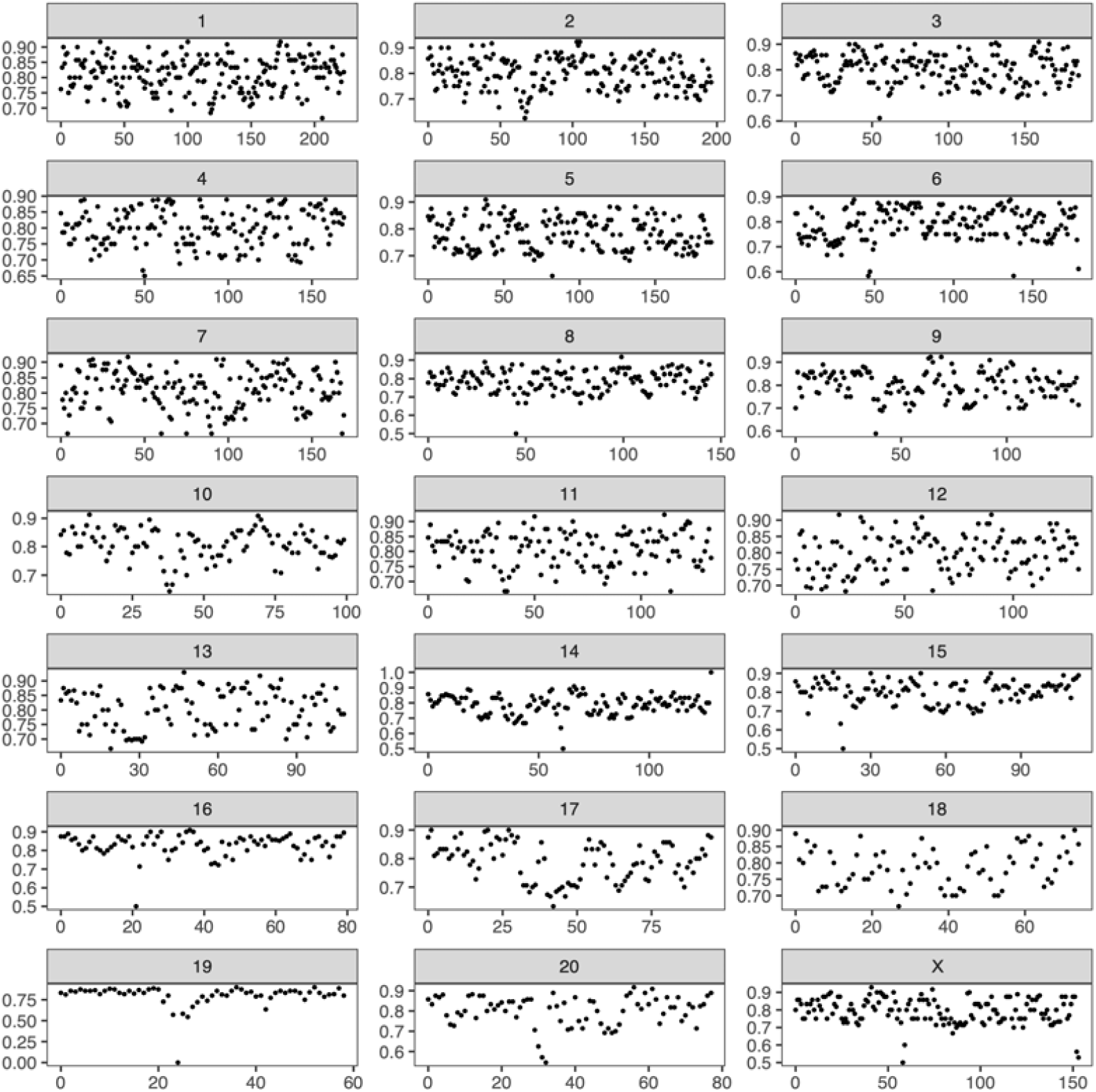
The chromosome-wide methylation frequencies for the Chinese rhesus macaque. The black points show the median values within slide-windows (the window-size of 1Mb).

**Figure 2.**
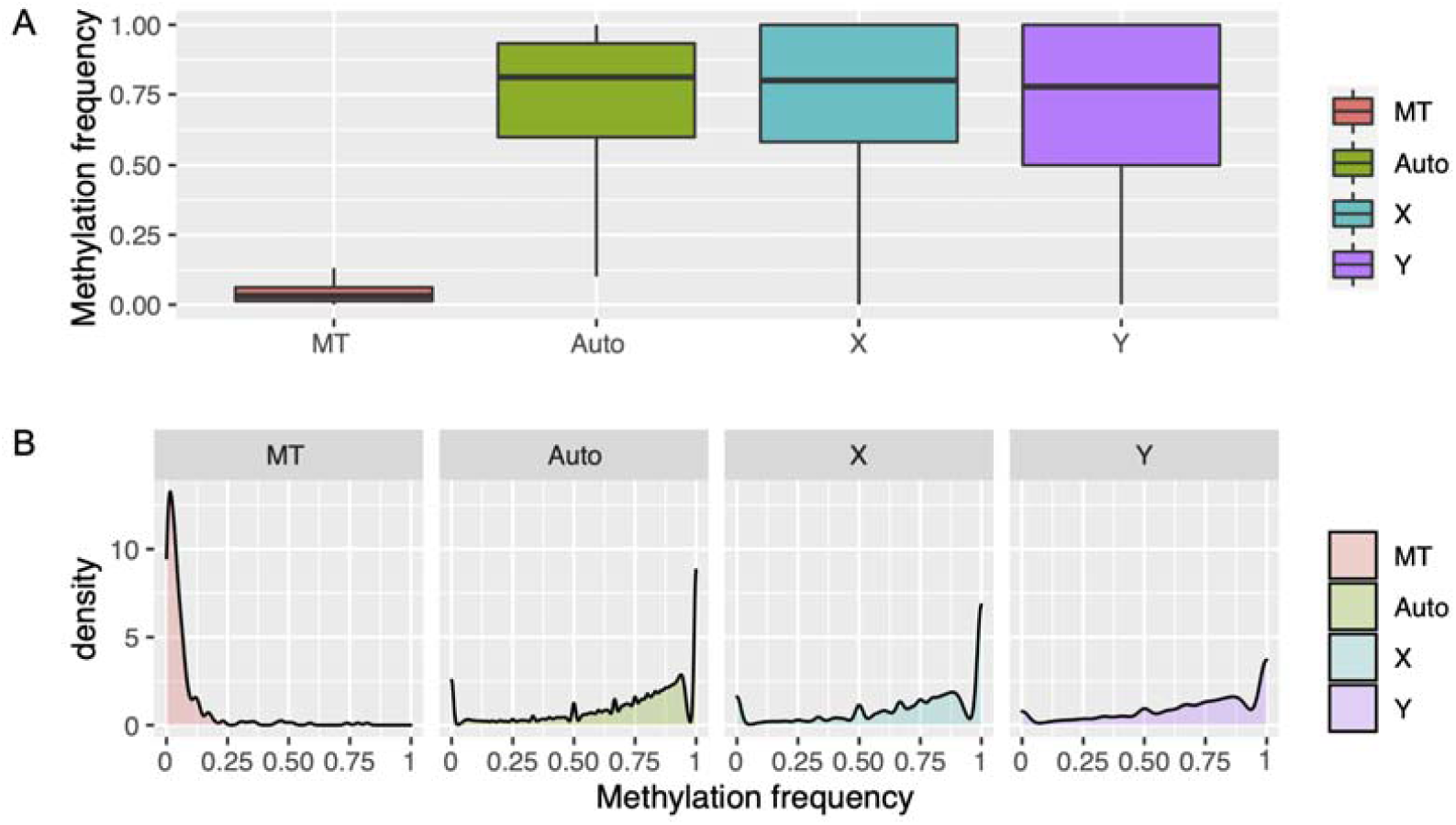
The distribution pattern of methylation frequencies across mitochondrial genome, autosomes, and sex chromosomes based on boxplot (A) and density plot (B). Only the “minimum (Q1 -1.5*IQR)”, first quartile [Q1], median, third quartile [Q3] and “maximum (Q3 + 1.5*IQR)” are shown for boxplot.

**Figure 3.**
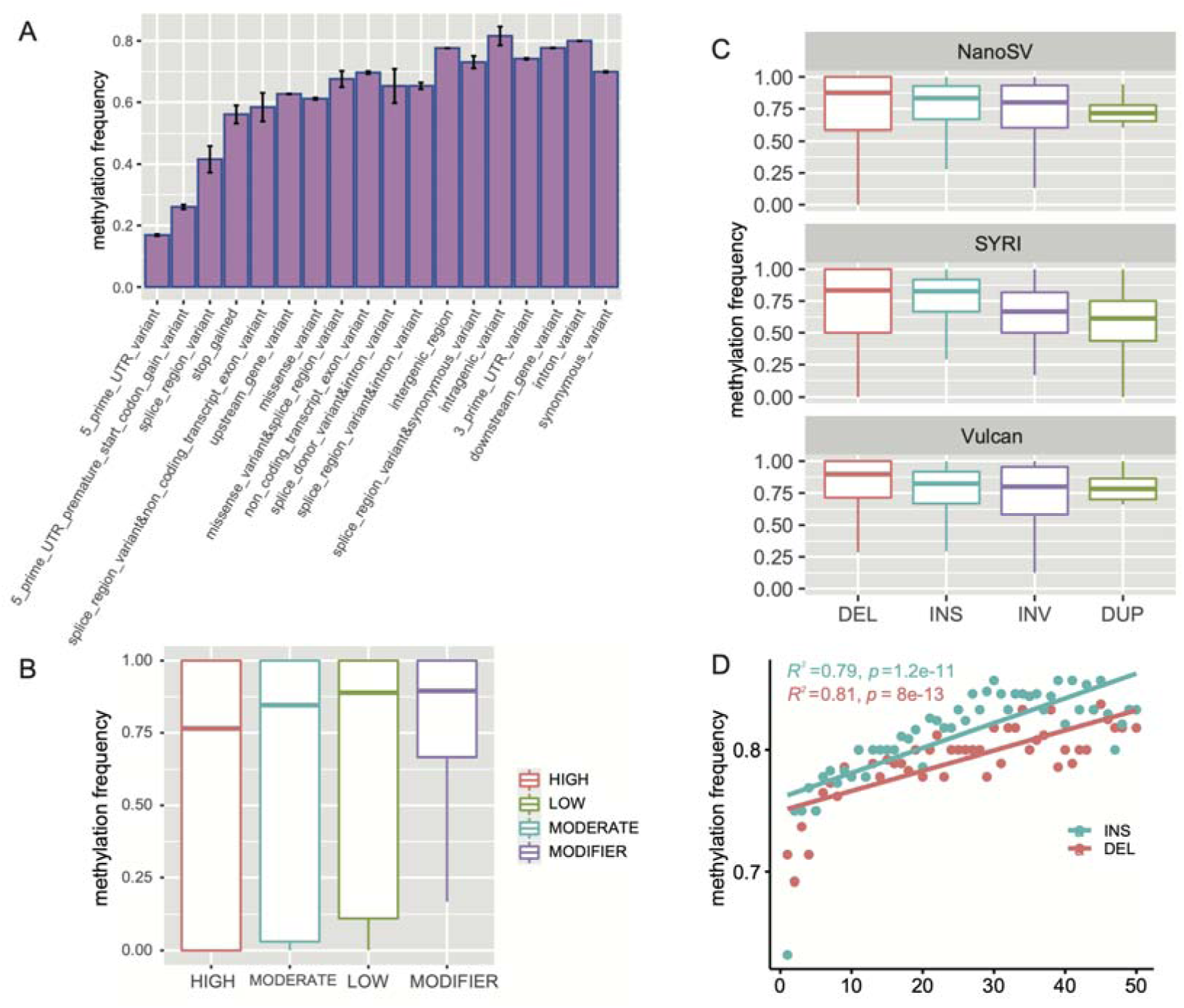
(A) The methylation frequencies of annotated SNPs within and around protein-coding genes. The tiny black bars show the standard errors. (B) Boxplots for methylation frequencies of SNPs with annotations of impacts on gene structure (following the annotation of SnpEff v5.1). (C) Boxplots for methylation frequencies of SVs from three different tools. (D) The median methylation frequencies across small Indels with different lengths. The formulas show linear regressions of median values against lengths of small Indels (< 50bp).

We comprehensively analyzed the methylation pattern for variants including SNPs, small indels (<50bp), and SVs (>50bp). We called variants of SNPs and small Insertion–deletion mutations (indels, <50bp) using the short-reads data of population genomes including 27 samples covering both Indian and Chinese subspecies. For SNPs, among variants annotated to affect different gene structures, we uncovered the lowest methylation levels for variants in 5’ UTR of genes (Figure 3A), suggesting a similar pattern with the previous finding that the methylation frequencies are the lowest around the transcription start sites (TSS) [65]. In addition, the SNPs predicted with high impact (e.g., transcript ablation, frameshift, etc.) showed a significant lower methylation frequency than those SNPs predicted with low (e.g., synonymous variants, etc.), modifier (e.g., 3’UTR region, 5’UTR region, etc.), and moderate (e.g., missense, 3’UTR deletion, etc.) impacts (Wilcoxon rank sum test, p = 2.31e-5, 5.80e-12, and 0.016, for low, modifier, and moderate impact SNPs, respectively) (Figure 3B).

For SVs called from three methods with exclusively long reads (NanoSV, Vulcan, and SyRI), we found similar patterns in methylation frequencies for different types of SVs (Figure 3C). Based on the distribution of median methylation frequencies for all four SVs types (deletions, 0.89; insertions, 0.83; inversions, 0.79; and duplications, 065), deletions and duplications were found to be the highest and the lowest methylated, respectively (Wilcoxon rank sum test, p < 0.05 for all pair-wise comparisons). For small indels, lengths of both deletions and insertions demonstrated significant positive correlations with methylation levels (Figure 3D), with deletions showing a higher positive correlation than insertions (0.81 vs. 0.79). In addition, methylation frequencies of small indels showed a significant lower median than those of SVs (0.80 vs. 0.83, Wilcoxon rank sum test, p = 0.016). These patterns suggested that variants impacting longer DNA segments may have higher levels of methylation.

### Duplications showed significant higher recombination rates than inversions, deletions, and insertions

It is long known that SVs are derived from the mutational process during meiosis by nonallelic homologous recombination (NAHR) [66]. However, it is still unclear whether recombination rates would be different among SVs and which type of SVs has the highest recombination rates. Here, we addressed this question by inferring the historical and long-term recombination rates (ρ/bp) using pyrho, which utilized population genomic data and took population demography into account [33]. This fine-scale and genome-wide mapping strategy revealed a similar landscape among chromosomes. Specifically, recombination rates were higher toward the distal regions of chromosomes and lower around the centromeres (Supplementary figure 3), which is consistent with the long-held knowledge of a “U shape” distribution at the chromosome level in animals and plants [67-70].

**Supplementary figure 3.**
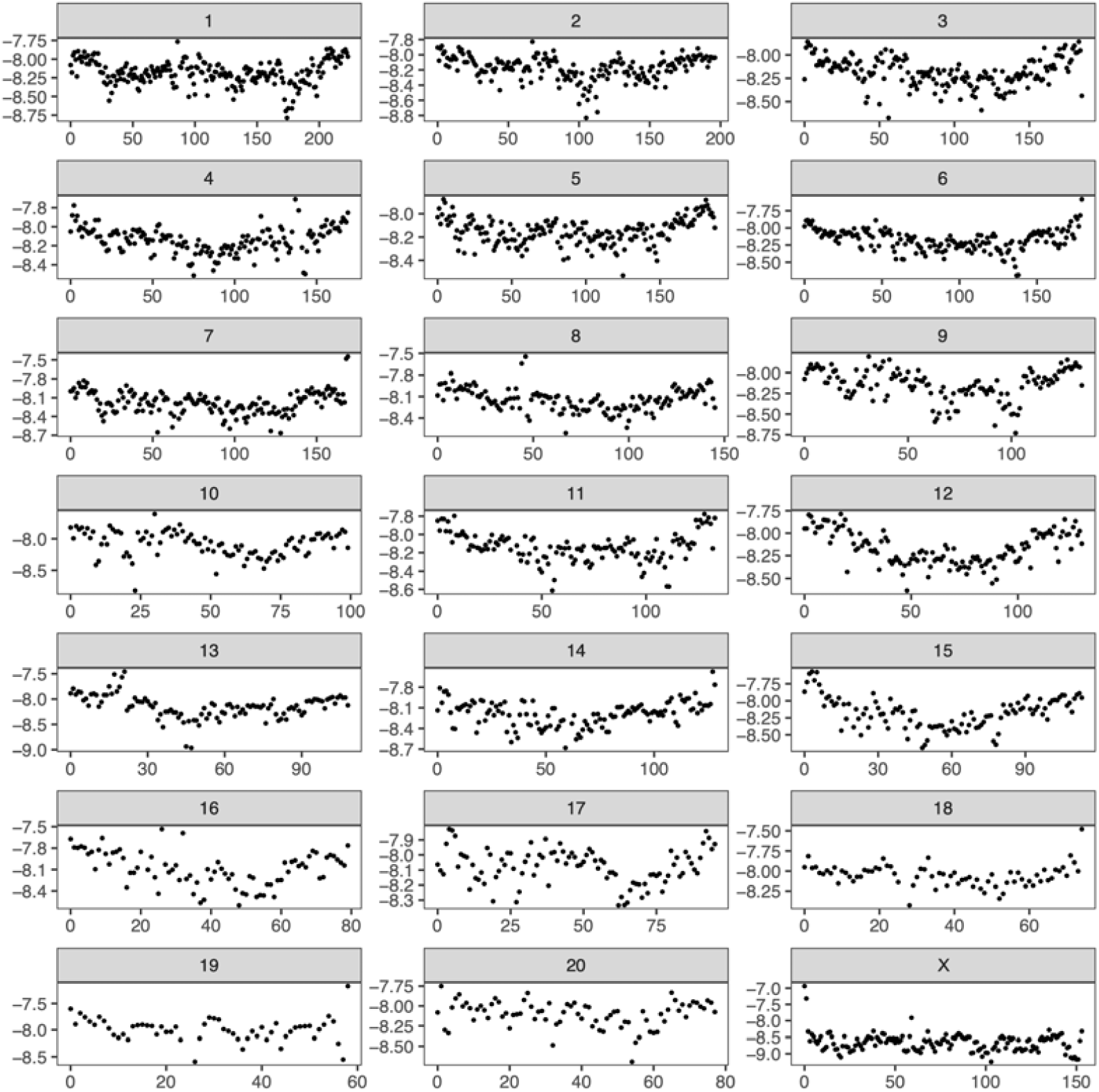
The fine-scale recombination rates inferred with pyrho package across chromosomes [33], which using population genomic data of the Chinese and Indian rhesus macaque (27 samples). The vertical axis is the logarithm of median fine-scale recombination rates with base 10 (ρ/Mb, slide-window, 1Mb). The horizontal axis shows the coordinates of chromosomes (Mb).

The comparison between four types of SVs called from three methods universally revealed that duplications showed significantly the highest fine-scale recombination rates (Wilcoxon rank sum test, p < 0.05 for all comparisons) (Figure 4A). This pattern was also consistent when comparing upstream and downstream regions (1000bp) of all SVs, suggesting no abrupt changes of fine-scale recombination rates within and around SVs. Interestingly, these comparisons were found to be more consistent for SVs identified from NanoSV and SyRI (Figure 4A). We further compared the distribution of recombination rates for four SVs types (Figure 4B). We found that the recombination rates of duplications showed two peaks, with one peak of recombination rates around 10^−8^, which is close to peaks of other three SVs types, and the other peak with higher recombination rate of 1.26×10^−7^.

**Figure 4.**
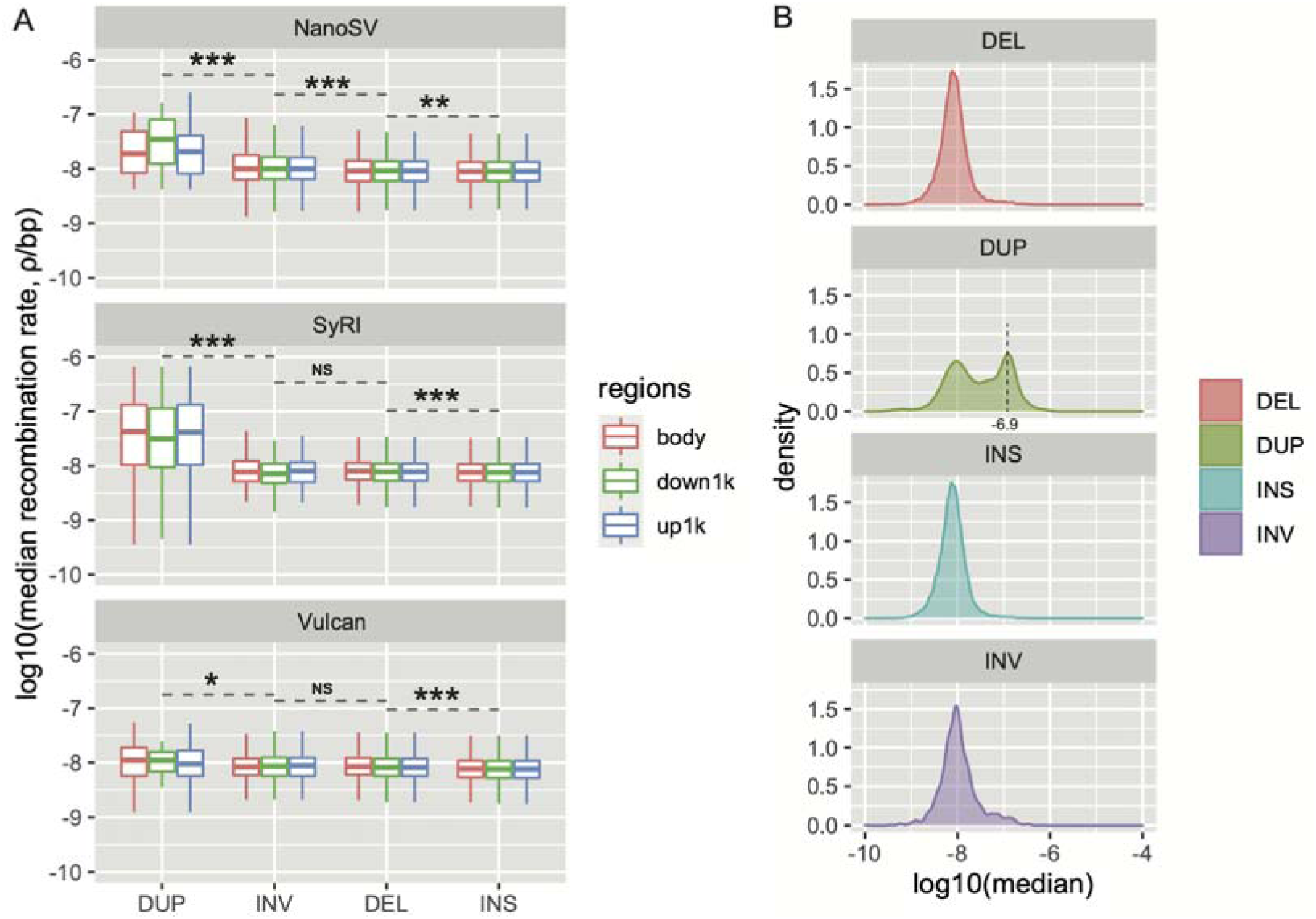
The distributions of common logarithm of recombination rates for SVs. (A) The boxplot comparisons for duplications, inversions, deletions, and insertions identified from three methods (NanoSV, SyRI, and Vulcan). The “body”, “down1k”, and “up1k” indicate the regions within the SVs, 1Kb downstream of the SVs, and 1Kb upstream of the SVs, respectively. “NS” indicates “non-significant” based on the Wilcoxon rank sum test (p>0.05). The significance levels of 0.05, 0.01, and 0.001 are showed with “*”, “**”, and “***”, respectively. (B) The density distribution of the fine-scale recombination rates for four types of SVs. The “INS”, “INV”, “DUP”, and “DEL” indicate the “insertions”, “inversions”, “duplications”, and “deletions”, respectively.

### X-chromosome contributes more SVs under positive selection

The population genetic theory predicts that positive Darwinian selection can drive the standing or *de novo* mutations to sweep through a population, to achieve diverse local adaptations and develop novel phenotypes [71]. To understand the influence of positive selection on the evolutionary fate of SVs, we inferred the regions under positive selection for the Chinese and Indian rhesus macaques. Based on population genetics, positive selection would hallmark several features along chromosomes, including but not limited to, the increased fixation index (*F*_*ST*_) between focal population and control population, the reduced nucleotide diversity (π) in focal population relative to control population, and the elevated expected haplotype homozygosity [72-74]. The selective sweep signals were estimated based on three complementary methods including the ratio of nucleotide diversity between Indian and Chinese populations (*θ*_π_India_/*θ*_π_China_), the fixation index (*F*_*ST*_), and the haplotype homozygosity (Ihh12) (Figure 5A-C).

**Figure 5.**
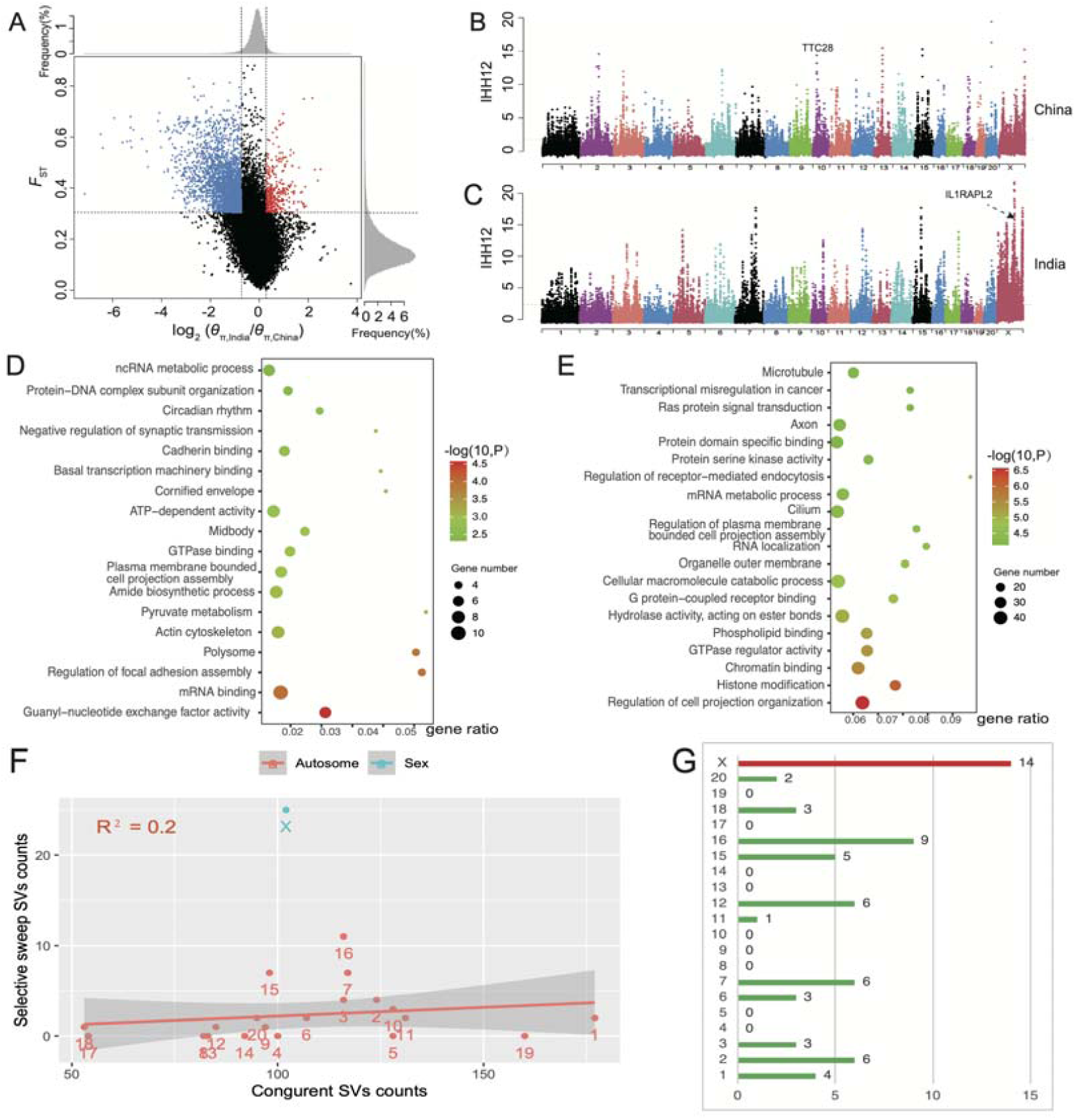
The selective signals for the Chinese and Indian rhesus macaques based on three complementary methods (F_st_, *θ*_π_India_/*θ*_π_China_, Ihh12, top 5% strongest windows, 100Kb). (A) The distribution of F_st_ and *θ*_π_India_/*θ*_π_China_ based on comparison between the Chinese and Indian rhesus macaques. The red and blue points show the the candidate selective sweep windows for the Indian and Chinese groups, respectively. (B) The candidate selective sweep signals above the dotted line (top 5% windows) for the Chinese rhesus macaques. The SVs involved gene (*TTC28*) with the highest signal in Ihh12 was shown. (C) The candidate selective sweep signals above the dotted line (top 5% windows) for the Indian rhesus macaques. The SVs involved gene (*IL1RAPL2*) with the highest signal in Ihh12 was shown. (D) The functional enrichment ontology for the Chinese rhesus populations. (E) The percentages of SVs under positive selection. (F) The chromosomal distribution for the 53 positively selected genes with signals of congruent SVs. Note: the horizontal axis shows the X chromosome while the vertical axis shows the number of genes. (G) The linear regression for congruent SVs counts and the SVs under selective sweep (p = 0.4, not significant). The blue point is the outlier (X chromosome).

Totally, we identified 1400 genes under selective sweep for the two subspecies, with 164 genes for the Chinese and 1236 genes for Indian rhesus macaques (Supplementary table 3). Notably, over 38% of positively selected genes were distributed in X chromosome, significantly higher than the expected average level of autosomes (the X^2^ test, p < 0.00001). The strongest signal in the Chinese group was found in an autosome gene *TTC28* (Figure 5B), which is related to human developmental disease based on human exome sequencing of fetal anomaly syndromes [75]. For Indian rhesus macaques, the highest signal was found in an X-chromosome gene, *IL1RAPL2* (Figure 5C), which is involved in diseases including mental development and autism [76, 77]. The gene function enrichment analysis revealed that the most significant pathways involve the Guanyl-nucleotide exchange factor activity and regulation of cell projection organization, for the Chinese (Figure 5D) and Indian rhesus macaques (Figure 5E), respectively. Interesting, we found that the two genes with the strongest signals among all positively selected genes, *IL1RAPL2* and *TTC28*, were involved in intragenic SVs (insertions and deletions, Supplementary table 4). Thus, it is very interesting to know whether SVs could be driven by the evolutionary force of positive selection in rhesus macaques. We found that, among the 2248 congruent SVs shared by three methods, 3.20% of SVs (72/2248) showed signals of selective sweep in intergenic and intragenic DNA regions (Supplementary table 5). Moreover, 3.79% of positively selected genes (53/1400) were detected with signals of congruent SVs (Supplementary table 6). Interestingly, X chromosome exhibited a significant higher contribution (26.45%, 14/53) to the positively selected genes involved in SVs than autosomes (the X^2^ test, p < 0.00001; Figure 5F). The regression analysis revealed the positively selected SVs in autosomes do not follow a simple linear model against the overall congruent SVs (p = 0.4, Figure 5F). However, the X chromosome exhibited as an obvious outlier with excess number of positively selected SVs (Figure 5G). These findings suggest that SVs could serve as an important raw material for natural selection to operate on genomic diversity.

## Discussion

One of core tenets in biology is that organisms adapt through natural selection on genomic variants to tackle ever-changing challenges. For evolutionary biologists of rhesus macaque, the genomic variants were of interest for addressing questions involving population demography [24], gene flow [78], disease variants [19], and natural selection [26]. At species level, rhesus macaque was reported to cover 2.5 times higher overall nucleotide diversity than humans, empowering this species as a great reservoir to discover and model corresponding pathogenic variants in human [19, 24]. At population level, the Chinese subspecies was found to host higher genomic diversity and higher effective population size (*Ne*) than the Indian subspecies [24, 26]. Thus, macaque species could serve as a powerful model to study intra-specific variants.

In medical and population genetics, structural variations (SVs, >50bp) are known with more pronounced phenotypic impacts than SNPs [79, 80]. Thus, SVs have increasingly become characterized in studies of human diseases including mental disorders and cancers [81]. However, as a critical medical model for human disease, the rhesus macaque was rarely investigated in issues related to the SVs. In this study, we initially presented a chromosome-level reference genome for a Chinese rhesus macaque (CR2) with both Nanopore long and Illumina short reads. Based on this reference and the nanopore long reads, we identified SVs between the Chinese and Indian subspecies and investigated their patterns and their relationship with methylation frequencies, recombination rates, and evolutionary forces.

We found that X chromosome is an obvious outlier for the significant linear pattern that shapes the number of SVs across autosomes. In addition, we detected significantly higher contribution of X-chromosomal SVs-involved genes to the reservoir of genes under selective sweep. These patterns are consistent with the well-known “faster-X effect”, which predicts that the X chromosome has a higher rate of adaptive evolution than the autosomes [63, 82, 83]. Although the “faster-X effect” is a generalization for the evolutionary divergence at the species level, our result suggests that the effect is more taxonomically pervasive than previously thought, especially at the subspecies level. Moreover, for medical genetics, the recognition of the “faster-X effect” at the population level could be informative to the dominance/recessivity of deleterious mutations leading to genetic diseases [28, 63, 84].

Taking advantages of nanopore data in providing information of DNA cytosine methylation [65], we estimated the methylation frequencies of the CR2 at a genome-wide scale. Previously, controversial conclusions were made for the methylation levels of mitochondrial genome relative to nuclear genome [85]. Very low or no methylation for mitochondrial genome was found by some studies [86, 87], although some contradictory conclusions were also made [88, 89]. Based on nanopore long reads, we found that mitochondrial genome has over 25 times lower median methylation frequency than nuclear chromosomes, thus providing evidence for the very low methylation level of mitochondrial genome. Interestingly, we found significantly lower methylation level for duplications than for other types of SVs (deletions, insertions, and inversions). Previous studies have found that the regions with decreased methylation tend to have higher GC content [90] and duplications demonstrate significantly higher GC content than non-duplications [91]. Thus, our finding of the decreased methylation level in duplications based on nanopore long reads is consistent with the implications of previous findings.

Duplications and other types of SVs are derived from the intergenerational mixing of DNA through homologous recombination during meiosis [92]. However, the relationship between recombination rates and different types of SVs is less clear. The yeast model has revealed that duplication is associated with the increased recombination rates [93]. In this study, we revealed significantly higher recombination rates in duplications than in other types of SVs. Half a century ago, Susumu Ohno has already proposed the notion of duplication as the major source of adaptive innovation during the long-term evolution [94]. Thus, the higher recombination rates may facilitate the evolution of duplications by boosting the efficiencies of purifying selection to remove deleterious duplications on the one hand, and positive selection to increase beneficial duplications on the other [95].

Consistent with the expectation of SVs as important raw materials upon which natural selection can then play, we found that over 3% of SVs may be under selective sweep. Moreover, over 3% of positively selected genes host signals of SVs. Some of these SVs-involved genes under positive selection have been reported with genetic defects related to human diseases. For example, the *SCN1A*, coding for the α-subunit of the neuronal voltage-gated sodium ion channel, type 1 (NaV 1.1), is correlated with human epileptic encephalopathies [96]. The genes *IL1RAPL2, ACSL4*, and *PAK3* are related to disease phenotypes including Alport syndrome, elliptocytosis, and intellectual disability [97, 98]. Interestingly, *SLC25A19*, a mitochondrial deoxyribonucleotide transporter, is related to severe congenital microcephaly (reduced brain size) [99-101]. From the perspectives of evolutionary biology and translational medicine, the SVs-involved genetic changes and the related positive selection in rhesus macaques could not only provide useful insights into the species- or subspecies-specific adaptation, but also facilitate the translational studies from nonhuman primate models to human. Altogether, our study and historical research may embark a promising journal to explore SVs-derived adaptive evolution and functional effects from the nonhuman primate models to our own species.

## Conclusion

Rhesus macaques (Macaca mulatta) is the most extensively used non-human primate medical disease model. The higher genomic diversity in this species than in human render it valuable for addressing medical as well as evolutionary questions. Despite the unique advantage of holding methylation signals, nanopore long-read data has not been utilized to investigate issues related to reference and variants. Here, we provided the long reads for a Chinese macaque (CR2) and conducted SVs-oriented analyses on genomic between two subspecies. We found lower strengths of methylation but higher rates of recombination for duplications than other types of SVs (deletions, insertions, and inversions). We also revealed the selective sweep and the “faster-X effect” underlying the SVs distribution along the chromosomes. Some SVs-involved genes under selective sweep are causal genes for multiple neurodevelopmental diseases and innovative traits in human. Our study characterizes multiple molecular features and evolutionary forces underlying SVs of the rhesus macaque, which could shed light on the translational medicine from this important medical model to human per se.

## Materials and methods

### DNA sampling, sequencing, and assembling

The genomic DNA was extracted from the blood sample of an adult Chinese rhesus macaque (male). The study protocol and data analyses were formally approved by the Ethics Committee of West China Hospital (Registration number: 20220211006). The long reads and short reads data were produced from Oxford PromethION and Illumina platform, respectively. The basecalling of Nanopore’s output was performed using Guppy [102] and only reads with mean quality scores >7 was retained.

To improve accuracy and continuity, we conducted *de novo* assembling based on the following three-step approach. Step one: long reads cleaning and *de novo* assembling. We conducted the first-round *de novo* assembling with NECAT v0.0.1 [103], which can correct errors of Nanopore long noisy reads before *de novo* assembling. The raw reads were corrected and then *de novo* assembled with default parameters except for “MIN_READ_LENGTH=1000”. Additionally, the NextDenovo v2.4.0 was also used for *de novo* assembly based on the cleaned reads of NECAT. We found a longer contig N50 for NextDenovo than for NECAT (65.99Mb vs. 6.44Mb). Thus, the contigs longer than 20Mb were used in step two. Step two: reference-guided chromosome assignment. To anchor the locations of contigs, we utilized a publicly available reference genome for the Chinese rhesus macaque (rheMacS) and the long contigs from NextDenovo as backbones to order and orient the NECAT raw contigs [34], with default parameters of the RagTag v2.1.0 [104]. Step three: SVs sensitive polishing. We combined several tools with default parameters to further polish the errors caused by long reads, including four rounds of racon v1.4.3 [105] with long reads and four rounds of pilon v1.24 [106] with short reads. We finally conducted misassemblies correction with tools sensitive to structural variations, including tigmint v1.2.6 [107] and Inspector v1.0.2 [108]. The final assembled genome was mapped to the Indian macaque reference (Mmul_10) with minimap2 [109] and visualized with dotplot by dotPlotly (https://github.com/tpoorten/dotPlotly).

Genome continuity was further evaluated with BUSCO metric [57] and QUAST statistics [110]. BUSCO can assess the completeness by the percentage of complete near-universal single-copy orthologs while QUAST statistics can be used to estimate N50, length distribution, etc. The genome was annotated with the long-read RNAseq data from a pool of tissues (heart, liver, whole brain, intestine, testis, muscle, and pancreas) using MAKER2 [111]. The “hidden genes” were retrieved from the Biomart tool from Ensembl (v105) with a focus on only the “one to one” orthologous genes between mouse, rat, human, and macaque. The genes that are only absent in macaque were determined as the “hidden genes”. These genes were then mapped back to the CR2 genome and only the genes with no mapping coordinates were defined as the real “hidden genes”. The recovery of these genes were conducted by mapping (BLAST tools, Evalue < 0.001) to the long-read RNAseq transcriptome obtained from the assembly-guided mapping and *de novo* assembly with tools including STAR v2.7.10a [112] and StringTie2 [113].

### Structural variations (SVs) identification and distribution

The SVs were identified using two complementary methods of “reads mapping” and “assembly comparison”. Specifically, for “reads-based” method, we explored two software, NanoSV [60] and Vulcan [61], to identify signals of SVs. NanoSV takes advantages of split- and gapped-aligned reads to define breakpoint-junctions of SVs, following the mapping of long reads to genome references (Mmul_10) with LAST v1256 [114] and the alignment processing with Sambamba V0.8.2 [115]. Vulcan integrates pipelines including dual mode alignment of long reads with aligners minimap2 [109] and NGMLR v0.2.7 [116] and SVs calling with Sniffles2 [116]. The “assembly-based” method was based on SyRI v1.6 [53]. We compared the results of SVs with BEDTools v2.30 [117]. The consensus SVs with shared regions (covering mutually at least 80% of SVs lengths) were identified as the lower bound of reliable call set. To reveal potentially consistent patterns of SVs from different algorithms, the SVs from these methods were compared and defined as the upper bound of reliable call set.

### Genome-wide methylation evaluation with nanopore sequencing data

The nanopore long-read data can be used to directly estimate epigenetic modifications at nucleotide resolution [31]. The signals of cytosine methylation were quantified with Nanopolish 0.14.0 [65]. The methylated sites and unmethylated sites were differentiated with log value of likelihood ratios > 2.5 and < -2.5, respectively [118].

### The inference of fine-scale recombination rates based on NGS population data

We inferred fine-scale recombination rates using pyrho [33] with population-wide NGS data, which include 27 samples from both Chinese and Indian subspecies. Different from the pedigree-based method, which can estimate the short-term signals of meiotic recombination, the fine-scale recombination rates take advantage of historical demography using the smc++ [119] and can recovery the population-wide and historical recombination at a genome-wide scale.

### The inference of selective sweep

The selective sweep was detected by combining the signals of the fixation index (*F*_*ST*_) between the Chinese and Indian rhesus macaque, the reduced nucleotide diversity (π) in Chinese rhesus population relative to Indian population (*θ*_π_India_/*θ*_π_China_), and the elevated expected haplotype homozygosity (Ihh12) [72-74]. The consensus of selective sweep regions was determined as the shared regions among the top 5% of empirical distribution of windows with the strongest signals for all three metrics.

## Supporting information

The tables for major results

## Acknowledgement

The authors report that no conflicts of interest exist. This study was supported by the fifth batch of technological innovation research projects in Chengdu (2021-YF05 -01331-SN), the Postdoctoral Research and Development Fund of West China Hospital of Sichuan University (2020HXBH087), and the Short-Term Expert Fund of West China Hospital (139190032). We also acknowledged the computing supports from the West China Biomedical Big Data Center and the Med-X Center for Informatics of Sichuan University.

## Author Contributions

B.R.S., C.Z.F., and G.Y. supervised this work. J.H.C., J.Z., Y.Z., Y.Y.J., and Y.N.C, designed the research. J.H.C., J.Z., Y.Y.J., and I. J. analyzed data. J.H.C. wrote the manuscript. I. J. and other authors helped to improve the manuscript.

## Competing interests

The authors declare no competing financial interests.

## Data availability

The raw data for the Chinese rhesus macaque (Macaca mulatta) used in this study can be retrieved from NCBI (ID: PRJNA889987, SAMN31264197).

## Notes

### Competing Interest Statement

The authors have declared no competing interest.

## References

1. Accogli, A., et al., Novel CNS malformations and skeletal anomalies in a patient with Beaulieu-boycott-Innes syndrome. Am J Med Genet A, 2018. 176(12): p. 2835–2840.

2. Gibbs, R.A., et al., Evolutionary and biomedical insights from the rhesus macaque genome. science, 2007. 316(5822): p. 222–234.

3. Van Rompay, K.K.A., Tackling HIV and AIDS: contributions by non-human primate models. Lab Animal, 2017. 46(6): p. 259–270.

4. Shedlock, D.J., G. Silvestri, and D.B. Weiner, Monkeying around with HIV vaccines: using rhesus macaques to define ‘gatekeepers’ for clinical trials. Nat Rev Immunol, 2009. 9(10): p. 717–28.

5. Geisbert, T.W., et al., Evaluation in nonhuman primates of vaccines against Ebola virus. Emerg Infect Dis, 2002. 8(5): p. 503–7.

6. Urano, E., et al., COVID-19 cynomolgus macaque model reflecting human COVID-19 pathological conditions. Proceedings of the National Academy of Sciences, 2021. 118(43): p. e2104847118.

7. He, W.-t., et al., Broadly neutralizing antibodies to SARS-related viruses can be readily induced in rhesus macaques. Science Translational Medicine, 2022. 14(657): p. eabl9605.

8. Cyranoski, D., Monkeys genetically modified to show autism symptoms. Nature, 2016. 529(7587): p. 449–449.

9. Zhao, H., Y.H. Jiang, and Y.Q. Zhang, Modeling autism in non-human primates: Opportunities and challenges. Autism Res, 2018. 11(5): p. 686–694.

10. Zhou, Y., et al., Atypical behaviour and connectivity in SHANK3-mutant macaques. Nature, 2019. 570(7761): p. 326–331.

11. Porras, G., Q. Li, and E. Bezard, Modeling Parkinson’s disease in primates: The MPTP model. Cold Spring Harb Perspect Med, 2012. 2(3): p. a009308.

12. Potts, L.F., et al., Modeling Parkinson’s disease in monkeys for translational studies, a critical analysis. Exp Neurol, 2014. 256: p. 133–43.

13. Miwa, Y., et al., Bilateral cataract surgery in a Japanese macaque (Macaca fuscata): A case report. Clin Case Rep, 2021. 9(11): p. e05112.

14. Lin, K.H., et al., Age-related changes in the rhesus macaque eye. Experimental Eye Research, 2021. 212: p. 108754.

15. Cox, L.A., et al., Nonhuman Primates and Translational Research-Cardiovascular Disease. Ilar j, 2017. 58(2): p. 235–250.

16. Tollner, T.L., et al., Macaque sperm coating protein DEFB126 facilitates sperm penetration of cervical mucus†. Human Reproduction, 2008. 23(11): p. 2523–2534.

17. Khampang, S., et al., Blastocyst development after fertilization with in&#xa0;vitro spermatids derived from nonhuman primate embryonic stem cells. F&S Science, 2021. 2(4): p. 365–375.

18. Cooper, E.B., et al., The rhesus macaque as a success story of the Anthropocene. Elife, 2022. 11.

19. Warren, W.C., et al., Sequence diversity analyses of an improved rhesus macaque genome enhance its biomedical utility. Science, 2020. 370(6523).

20. Weiss, A., et al., Rhesus macaques (Macaca mulatta) as living fossils of hominoid personality and subjective well-being. J Comp Psychol, 2011. 125(1): p. 72–83.

21. Wu, S.-J., et al., Ecological Genetics of Chinese Rhesus Macaque in Response to Mountain Building: All Things Are Not Equal. PLOS ONE, 2013. 8(2): p. e55315.

22. Hasan, M.K., et al., Distribution of Rhesus Macaques (Macaca mulatta) in Bangladesh: Inter-population Variation in Group Size and Composition. Primate Conservation, 2013. 26(1): p. 125-132, 8.

23. Hernandez, R.D., et al., Demographic histories and patterns of linkage disequilibrium in Chinese and Indian rhesus macaques. Science, 2007. 316(5822): p. 240–243.

24. Xue, C., et al., The population genomics of rhesus macaques (Macaca mulatta) based on whole-genome sequences. Genome research, 2016. 26(12): p. 1651–1662.

25. Bimber, B.N., et al., mGAP: the macaque genotype and phenotype resource, a framework for accessing and interpreting macaque variant data, and identifying new models of human disease. BMC Genomics, 2019. 20(1): p. 176.

26. Liu, Z., et al., Population genomics of wild Chinese rhesus macaques reveals a dynamic demographic history and local adaptation, with implications for biomedical research. GigaScience, 2018. 7(9).

27. Chen, J., et al., The de novo assembly of a European wild boar genome revealed unique patterns of chromosomal structural variations and segmental duplications. Animal Genetics, 2022. 53(3): p. 281–292.

28. Vicoso, B. and B. Charlesworth, Evolution on the X chromosome: unusual patterns and processes. Nature Reviews Genetics, 2006. 7(8): p. 645–653.

29. Villegas-Mirón, P., et al., Chromosome X-wide Analysis of Positive Selection in Human Populations: Common and Private Signals of Selection and its Impact on Inactivated Genes and Enhancers. Frontiers in Genetics, 2021. 12.

30. Bechsgaard, J., et al., Evidence for Faster X Chromosome Evolution in Spiders. Molecular Biology and Evolution, 2019. 36(6): p. 1281–1293.

31. Liu, Y., et al., DNA methylation-calling tools for Oxford Nanopore sequencing: a survey and human epigenome-wide evaluation. Genome Biology, 2021. 22(1): p. 295.

32. Akbari, V., et al., Genome-wide detection of imprinted differentially methylated regions using nanopore sequencing. Elife, 2022. 11.

33. Spence, J.P. and Y.S. Song, Inference and analysis of population-specific fine-scale recombination maps across 26 diverse human populations. Science Advances, 2019. 5(10): p. eaaw9206.

34. He, Y., et al., Long-read assembly of the Chinese rhesus macaque genome and identification of ape-specific structural variants. Nature Communications, 2019. 10(1): p. 4233.

35. Alhakami, H., H. Mirebrahim, and S. Lonardi, A comparative evaluation of genome assembly reconciliation tools. Genome Biol, 2017. 18(1): p. 93.

36. Deamer, D., M. Akeson, and D. Branton, Three decades of nanopore sequencing. Nature biotechnology, 2016. 34(5): p. 518–524.

37. Loman, N.J., J. Quick, and J.T. Simpson, A complete bacterial genome assembled de novo using only nanopore sequencing data. Nature methods, 2015. 12(8): p. 733–735.

38. Jain, M., et al., Nanopore sequencing and assembly of a human genome with ultra-long reads. Nature Biotechnology, 2018. 36(4): p. 338–345.

39. Xie, H., et al., De novo assembly of human genome at single-cell levels. Nucleic Acids Research, 2022. 50(13): p. 7479–7492.

40. Nurk, S., et al., The complete sequence of a human genome. Science, 2022. 376(6588): p. 44–53.

41. Jain, M., et al., Nanopore sequencing and assembly of a human genome with ultra-long reads. Nature biotechnology, 2018. 36(4): p. 338–345.

42. Edwards, R.J., et al., Chromosome-length genome assembly and structural variations of the primal Basenji dog (Canis lupus familiaris) genome. BMC Genomics, 2021. 22(1): p. 188.

43. Chueca, L.J., et al., De novo Genome Assembly of the Raccoon Dog (Nyctereutes procyonoides). Frontiers in Genetics, 2021. 12.

44. Wang, C., et al., A novel canine reference genome resolves genomic architecture and uncovers transcript complexity. Communications Biology, 2021. 4(1): p. 185.

45. Dhar, R., et al., De novo assembly of the Indian blue peacock (Pavo cristatus) genome using Oxford Nanopore technology and Illumina sequencing. GigaScience, 2019. 8(5).

46. Warren, W.C., et al., A New Chicken Genome Assembly Provides Insight into Avian Genome Structure. G3 Genes|Genomes|Genetics, 2017. 7(1): p. 109–117.

47. Choi, J.Y., et al., Nanopore sequencing-based genome assembly and evolutionary genomics of circum-basmati rice. Genome Biology, 2020. 21(1): p. 21.

48. Tanaka, T., et al., De novo Genome Assembly of the indica Rice Variety IR64 Using Linked-Read Sequencing and Nanopore Sequencing. G3 (Bethesda), 2020. 10(5): p. 1495–1501.

49. Wu, T., et al., The De Novo Genome Assembly of Olea europaea subsp. cuspidate, a Widely Distributed Olive Close Relative. Frontiers in Genetics, 2022. 13.

50. Song, D., et al., Chromosome-Level Genome Assembly of the Burbot (Lota lota) Using Nanopore and Hi-C Technologies. Frontiers in Genetics, 2021. 12.

51. Sun, L., et al., Chromosome-level genome assembly of a cyprinid fish Onychostoma macrolepis by integration of nanopore sequencing, Bionano and Hi-C technology. Molecular Ecology Resources, 2020. 20(5): p. 1361–1371.

52. Kim, B.Y., et al., Highly contiguous assemblies of 101 drosophilid genomes. eLife, 2021. 10: p. e66405.

53. Goel, M., et al., SyRI: finding genomic rearrangements and local sequence differences from whole-genome assemblies. Genome Biology, 2019. 20(1): p. 277.

54. Liu, B., et al., Estimation of genomic characteristics by analyzing k-mer frequency in de novo genome projects. arXiv preprint 1308.2012, 2013.

55. Gavrielatos, M., et al., Benchmarking of next and third generation sequencing technologies and their associated algorithms for de novo genome assembly. Molecular Medicine Reports, 2021. 23(4): p. 1–1.

56. Johnson, L.K., et al., Draft genome assemblies using sequencing reads from Oxford Nanopore Technology and Illumina platforms for four species of North American Fundulus killifish. Gigascience, 2020. 9(6): p. giaa067.

57. Manni, M., et al., BUSCO: Assessing Genomic Data Quality and Beyond. Current Protocols, 2021. 1(12): p. e323.

58. Soneson, C., et al., A comprehensive examination of Nanopore native RNA sequencing for characterization of complex transcriptomes. Nature Communications, 2019. 10(1): p. 3359.

59. van Belzen, I.A.E.M., et al., Structural variant detection in cancer genomes: computational challenges and perspectives for precision oncology. npj Precision Oncology, 2021. 5(1): p. 15.

60. Cretu Stancu, M., et al., Mapping and phasing of structural variation in patient genomes using nanopore sequencing. Nature Communications, 2017. 8(1): p. 1326.

61. Fu, Y., et al., Vulcan: Improved long-read mapping and structural variant calling via dual-mode alignment. GigaScience, 2021. 10(9).

62. Wu, Z., et al., Structural variants in the Chinese population and their impact on phenotypes, diseases and population adaptation. Nature Communications, 2021. 12(1): p. 6501.

63. Meisel, R.P. and T. Connallon, The faster-X effect: integrating theory and data. Trends in Genetics, 2013. 29(9): p. 537–544.

64. Schatz, M.C., Nanopore sequencing meets epigenetics. Nature Methods, 2017. 14(4): p. 347–348.

65. Simpson, J.T., et al., Detecting DNA cytosine methylation using nanopore sequencing. Nature Methods, 2017. 14(4): p. 407–410.

66. Stankiewicz, P. and J.R. Lupski, Genome architecture, rearrangements and genomic disorders. Trends in Genetics, 2002. 18(2): p. 74–82.

67. Singhal, S., et al., Stable recombination hotspots in birds. Science, 2015. 350(6263): p. 928–932.

68. Tortereau, F., et al., A high density recombination map of the pig reveals a correlation between sex-specific recombination and GC content. BMC Genomics, 2012. 13(1): p. 586.

69. Dreissig, S., et al., Natural variation in meiotic recombination rate shapes introgression patterns in intraspecific hybrids between wild and domesticated barley. New Phytologist, 2020. 228(6): p. 1852–1863.

70. Huang, K., et al., Mutation Load in Sunflower Inversions Is Negatively Correlated with Inversion Heterozygosity. Molecular Biology and Evolution, 2022. 39(5).

71. Pritchard, J.K., J.K. Pickrell, and G. Coop, The genetics of human adaptation: hard sweeps, soft sweeps, and polygenic adaptation. Curr Biol, 2010. 20(4): p. R208–15.

72. Harris, A.M., N.R. Garud, and M. DeGiorgio, Detection and Classification of Hard and Soft Sweeps from Unphased Genotypes by Multilocus Genotype Identity. Genetics, 2018. 210(4): p. 1429–1452.

73. Porto-Neto, L.R., et al., Detection of Signatures of Selection Using FST, in Genome-Wide Association Studies and Genomic Prediction, xC. Gondro, J. van der Werf, and B. Hayes, Editors. 2013, Humana Press: Totowa, NJ. p. 423–436.

74. James, J., D. Castellano, and A. Eyre-Walker, DNA sequence diversity and the efficiency of natural selection in animal mitochondrial DNA. Heredity, 2017. 118(1): p. 88–95.

75. Meier, N., et al., Exome sequencing of fetal anomaly syndromes: novel phenotype–genotype discoveries. European Journal of Human Genetics, 2019. 27(5): p. 730–737.

76. Zhang, K.-J., et al., An association study between IL1RAPL2 gene and non-specific mental retardation in Chinese children. Genes & Genomics, 2010. 32(2): p. 159–164.

77. Kantojärvi, K., et al., Fine mapping of Xq11. 1-q21. 33 and mutation screening of RPS6KA6, ZNF711, ACSL4, DLG3, and IL1RAPL2 for autism spectrum disorders (ASD). Autism Research, 2011. 4(3): p. 228–233.

78. Evans, B.J., et al., Speciation over the edge: gene flow among non-human primate species across a formidable biogeographic barrier. Royal Society Open Science, 2017. 4(10): p. 170351.

79. Chen, L., et al., Association of structural variation with cardiometabolic traits in Finns. The American Journal of Human Genetics, 2021. 108(4): p. 583–596.

80. Collins, R.L., et al., A structural variation reference for medical and population genetics. Nature, 2020. 581(7809): p. 444–451.

81. Stankiewicz, P. and J.R. Lupski, Structural variation in the human genome and its role in disease. Annu Rev Med, 2010. 61: p. 437–55.

82. Charlesworth, B., J.A. Coyne, and N.H. Barton, The Relative Rates of Evolution of Sex Chromosomes and Autosomes. The American Naturalist, 1987. 130(1): p. 113–146.

83. Mank, J.E., et al., EFFECTIVE POPULATION SIZE AND THE FASTER-X EFFECT: EMPIRICAL RESULTS AND THEIR INTERPRETATION. Evolution, 2010. 64(3): p. 663–674.

84. Ellegren, H. and J. Parsch, The evolution of sex-biased genes and sex-biased gene expression. Nat Rev Genet, 2007. 8(9): p. 689–98.

85. Guitton, R., G.S. Nido, and C. Tzoulis, No evidence of extensive non-CpG methylation in mtDNA. Nucleic Acids Research, 2022. 50(16): p. 9190–9194.

86. Hong, E.E., et al., Regionally Specific and Genome-Wide Analyses Conclusively Demonstrate the Absence of CpG Methylation in Human Mitochondrial DNA. Molecular and Cellular Biology, 2013. 33(14): p. 2683–2690.

87. Maekawa, M., et al., Methylation of Mitochondrial DNA Is Not a Useful Marker for Cancer Detection. Clinical Chemistry, 2004. 50(8): p. 1480–1481.

88. Bianchessi, V., et al., Methylation profiling by bisulfite sequencing analysis of the mtDNA Non-Coding Region in replicative and senescent Endothelial Cells. Mitochondrion, 2016. 27: p. 40–47.

89. Feng, S., et al., Correlation between increased ND2 expression and demethylated displacement loop of mtDNA in colorectal cancer. Molecular medicine reports, 2012. 6(1): p. 125–130.

90. Tang, C.S.M. and R.J. Epstein, A Structural Split in the Human Genome. PLOS ONE, 2007. 2(7): p. e603.

91. Nakken, S., et al., Large-scale inference of the point mutational spectrum in human segmental duplications. BMC Genomics, 2009. 10(1): p. 43.

92. Kong, A., et al., Recombination rate and reproductive success in humans. Nature genetics, 2004. 36(11): p. 1203–1206.

93. Fang, O., et al., Genome Duplication Increases Meiotic Recombination Frequency: A Saccharomyces cerevisiae Model. Molecular biology and evolution, 2021. 38(3): p. 777–787.

94. Ohno, S., Evolution by gene duplication. 1970: Springer Science & Business Media.

95. Dapper, A.L. and B.A. Payseur, Connecting theory and data to understand recombination rate evolution. Philosophical Transactions of the Royal Society B: Biological Sciences, 2017. 372(1736): p. 20160469.

96. Parihar, R. and S. Ganesh, The SCN1A gene variants and epileptic encephalopathies. Journal of Human Genetics, 2013. 58(9): p. 573–580.

97. Piccini, M., et al., FACL4, a New Gene Encoding Long-Chain Acyl-CoA Synthetase 4, Is Deleted in a Family with Alport Syndrome, Elliptocytosis, and Mental Retardation. Genomics, 1998. 47(3): p. 350–358.

98. Allen, K.M., et al., PAK3 mutation in nonsyndromic X-linked mental retardation. Nature Genetics, 1998. 20(1): p. 25–30.

99. Lindhurst, M.J., et al., Knockout of Slc25a19 causes mitochondrial thiamine pyrophosphate depletion, embryonic lethality, CNS malformations, and anemia. Proceedings of the National Academy of Sciences, 2006. 103(43): p. 15927–15932.

100. Spiegel, R., et al., SLC25A19 mutation as a cause of neuropathy and bilateral striatal necrosis. Annals of Neurology, 2009. 66(3): p. 419–424.

101. Bottega, R., et al., Functional analysis of the third identified SLC25A19 mutation causative for the thiamine metabolism dysfunction syndrome 4. Journal of Human Genetics, 2019. 64(11): p. 1075–1081.

102. Wick, R.R., L.M. Judd, and K.E. Holt, Performance of neural network basecalling tools for Oxford Nanopore sequencing. Genome Biology, 2019. 20(1): p. 129.

103. Chen, Y., et al., Efficient assembly of nanopore reads via highly accurate and intact error correction. Nature Communications, 2021. 12(1): p. 60.

104. Alonge, M., et al., RaGOO: fast and accurate reference-guided scaffolding of draft genomes. Genome biology, 2019. 20(1): p. 1–17.

105. Vaser, R., et al., Fast and accurate de novo genome assembly from long uncorrected reads. Genome Res, 2017. 27(5): p. 737–746.

106. Walker, B.J., et al., Pilon: An Integrated Tool for Comprehensive Microbial Variant Detection and Genome Assembly Improvement. PLOS ONE, 2014. 9(11): p. e112963.

107. Jackman, S.D., et al., Tigmint: correcting assembly errors using linked reads from large molecules. BMC Bioinformatics, 2018. 19(1): p. 393.

108. Chen, Y., et al., Accurate long-read de novo assembly evaluation with Inspector. Genome Biology, 2021. 22(1): p. 312.

109. Li, H., Minimap2: pairwise alignment for nucleotide sequences. Bioinformatics, 2018. 34(18): p. 3094–3100.

110. Gurevich, A., et al., QUAST: quality assessment tool for genome assemblies. Bioinformatics, 2013. 29(8): p. 1072–1075.

111. Cantarel, B.L., et al., MAKER: an easy-to-use annotation pipeline designed for emerging model organism genomes. Genome research, 2008. 18(1): p. 188–196.

112. Dobin, A., et al., STAR: ultrafast universal RNA-seq aligner. Bioinformatics, 2013. 29(1): p. 15–21.

113. Kovaka, S., et al., Transcriptome assembly from long-read RNA-seq alignments with StringTie2. Genome Biology, 2019. 20(1): p. 278.

114. Kielbasa, S.M., et al., Adaptive seeds tame genomic sequence comparison. Genome Research, 2011. 21(3): p. 487–493.

115. Tarasov, A., et al., Sambamba: fast processing of NGS alignment formats. Bioinformatics, 2015. 31(12): p. 2032–2034.

116. Sedlazeck, F.J., et al., Accurate detection of complex structural variations using single-molecule sequencing. Nature Methods, 2018. 15(6): p. 461–468.

117. Quinlan, A.R. and I.M. Hall, BEDTools: a flexible suite of utilities for comparing genomic features. Bioinformatics, 2010. 26(6): p. 841–842.

118. Greenberg, M.V. and D. Bourc’his, The diverse roles of DNA methylation in mammalian development and disease. Nature reviews Molecular cell biology, 2019. 20(10): p. 590–607.

119. Steinrücken, M., et al., Inference of complex population histories using whole-genome sequences from multiple populations. Proceedings of the National Academy of Sciences, 2019. 116(34): p. 17115–17120.

